# Understanding Foundation Models in Digital Pathology: Performance, Trade-offs, and Model-Selection Recommendations

**DOI:** 10.1101/2025.09.12.675923

**Authors:** Danial Maleki, Nazim Shaikh, Xiao Li, Yao Nie, Raghavan Venugopal, Uday Kurkure

## Abstract

The rapid proliferation of digital pathology foundation models (FMs), spanning widely in architectural scales and pre-training datasets, poses a significant challenge in selecting the optimal model for a specific clinical application. To systematically evaluate these performance trade-offs, we conducted a comprehensive benchmark of five FMs, stratified from small to huge scales, across a diverse suite of whole slide image (WSI) and region of interest (ROI) tasks. Our findings demonstrate that model superiority is strongly task-specific, challenging the assumption that a larger scale universally offers an advantage. For instance, while the huge-scale Virchow2 model excelled at WSI metastasis detection, the small-scale Lunit model was superior for fine-grained lung subtype classification. This trend was even more pronounced in survival analysis, where smaller models outperformed their massive-scale counterparts, a phenomenon we hypothesize is linked to potential information bottlenecks within downstream aggregation models. Notably, the base-scale Kaiko model consistently provided a compelling balance, delivering competitive accuracy, superior stability in prognostic tasks, and higher computational efficiency. Our analysis suggests that the optimal FM is not necessarily the largest, but one whose scale, data composition, and training strategy are best aligned with the specific task. This work offers a practical, evidence-based framework for balancing performance, stability, and real-world deployment costs in computational pathology.

## Introduction

Digital Pathology foundation models (FMs) are large-scale deep learning systems designed to learn transferable visual representations from vast repositories of histopathology images. By pretraining directly on millions of whole-slide images (WSIs) in a self-supervised (learning from the image data itself) or weakly supervised manner (e.g., using slide-level diagnostic labels from pathology reports), these models capture morphological patterns across tissue types, staining variations, and disease contexts. The resulting feature extractors consistently outperform^1^ conventional supervised models or vision backbones pretrained on natural image datasets such as ImageNet.

Evidence from recent studies demonstrates that Digital Pathology FMs achieve state-of-the-art performance across diverse diagnostic and prognostic tasks^2^. In diagnosis, these models excel at identifying the fundamental characteristics of a disease from routine H&E slides. This includes precise tumor subtype classification, the detection of rare histological variants, and the prediction of underlying molecular features directly from morphology^1^. For instance, FMs can predict genomic alterations like microsatellite instability (MSI) or assess protein expression levels for biomarkers such as HER2 and PD-L1, potentially reducing the need for ancillary immunohistochemistry or molecular assays. In prognosis, FMs can forecast disease progression and patient futures. By analyzing morphological patterns invisible to the human eye, they provide powerful survival risk stratification^3^, helping to predict patient outcomes more accurately. These advances highlight the potential integration of FMs into clinical decision-support pipelines, enabling rapid, cost-effective, and scalable pathology workflows.

As the field has matured, the ecosystem of available FMs has expanded rapidly in both model diversity and scale (see Section). New models differ significantly in their network architecture (e.g., Convolutional Neural Networks (CNN), Vision Transformer (ViT) - based, or hybrid designs), pretraining objectives (e.g., contrastive learning, masked image modelling), and crucially — the scale and diversity of their training datasets. This proliferation has made it increasingly important for researchers and clinicians to understand the relative strengths, and weaknesses of each model to select the most suitable one for their specific research or diagnostic tasks.

Recent large-scale benchmarking studies have begun to map this complex landscape, each focusing on a different dimension of model performance and utility. For instance,^4^ benchmarked four foundation models to evaluate various adaptation strategies, testing different fine-tuning methods and using few-shot learning to gauge adaptability in data-limited environments.^1^ presented a clinical benchmark using datasets from standard hospital operations to assess models on clinically relevant tasks, analyzing performance against model size and inference costs. In another comprehensive study,^5^ evaluated 19 foundation models on “truly external” cohorts to avoid data leakage, finding that the diversity of pretraining data often outweighs sheer data volume. While this foundational work provides critical insights, it also reveals a practical gap — the direct cost-benefit implications of model scale remain poorly understood. It remains unclear how performance trade-offs between smaller and larger models manifest across diverse tasks, or how the scale of pretraining data interacts with model size. To address this gap, our work provides an evidence-based, practically oriented benchmark focused on model scale. We aim to answer the following key questions:

1. How does the scale of a foundation model directly impact its performance across a diverse range of downstream pathology tasks?
2. Are models trained on smaller datasets sufficient to achieve strong downstream performance, or do we need larger models trained on larger datasets, in order to fully realize their potentials on downstream tasks?
3. Is model superiority task-specific, or do models that excel at one task tend to generalize their high performance across others?

### Related work

he landscape of computational pathology has been reshaped by the rise of self-supervised learning (SSL) and the adoption of Vision Transformers (ViTs) as the de facto architecture for foundation models (FMs). This paradigm shift has spurred a wave of innovation, leading to a diverse ecosystem of models. Architectural developments range from hybrid designs like CTransPath^6^ that integrate convolutional layers with a Swin Transformer, to specialized lightweight models like PathDino^7^, and novel architectures such as Prov-GigaPath^8^, which adapts the LongNet method for whole-slide-level reasoning. Alongside this architectural evolution, the field is characterized by a rapid escalation in scale. FMs now span a vast range of sizes, from efficient models with only 9 million parameters^7^, trained on only 11 thousand WSIs to giant-scale systems like Virchow2^9^ with nearly two billion parameters and trained on millions of WSIs. Figure 1 provides a comparative visualization of these models with respect to model size and the number of WSIs used during training. To better understand this complex and rapidly evolving landscape, it is useful to categorize these models by their architectural scale and computational requirements.

**Figure 1.**
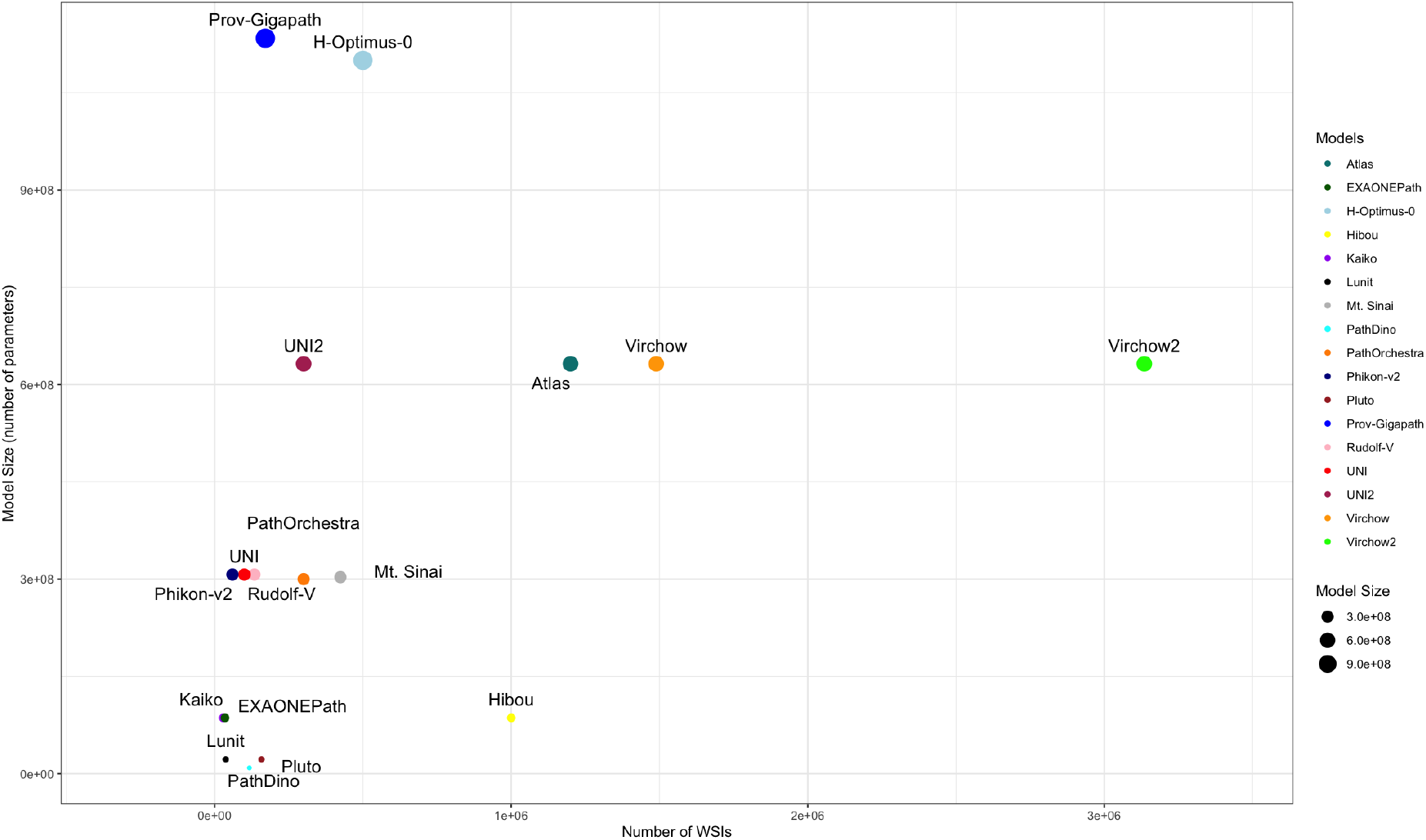
Comparative landscape of Digital Pathology FMs, organized by model size (number of parameters) and their training data size (number of WSIs).

#### The Spectrum of Digital Pathology Foundation Models

- *Small-Scale Models*: These models typically have fewer than 22 million parameters. Examples include Lunit’s^10^ DINO variant, the ViT-Small variant of Kaiko^11^, and PLUTO^12^. These models are characterized by their minimal computational requirements, making them ideal for efficiency-focused benchmarks.
- *Base-Scale Models:* Models in this tier generally have around 86 million parameters. Examples include the ViT-Base version of Kaiko^11^ and EXAONEPath^13^. They often can serve as alternatives for small-scale models in scenarios requiring higher performance while meeting deployment constraints, representing a sweet spot between strong feature representation and computational cost.
- *Large-Scale Models:* This category includes models with around 300 million parameters such as Phikon-v2^14^ and RudolfV^15^. These models often leverage advanced pre-training objectives to generate highly descriptive and generalizable representations, suitable for complex research problems involving diverse datasets.
- *Huge-Scale and Giant-Scale Models:* Sitting at the upper end of the spectrum, these models have parameters ranging from 632 million to over a billion. Key examples include UNI2^16^, Virchow2^9^, the billion-parameter H-Optimus-0^17^ and Prov-GigaPath^8^ models. By leveraging massive WSI training corpora, coupled with larger architectures and sophisticated training objectives, these models may have capability to achieve strong pan-cancer transferability and cross-domain robustness.

#### Contributions of this study

This study investigates the practical trade-offs of foundation model adoption.Moving beyond general performance benchmarks, We address which models are most appropriate for specific contexts defined by computational budgets, data availability, and performance expectations. To answer this, our work makes several key contributions

1. We select and benchmark a stratified set of five state-of-the-art models that represent four distinct tiers of scale:
  a. *Lunit:* ViT-Small model pretrained on the 29K TCGA^18^ dataset.
  b. *Kaiko:* ViT-Base model also pretrained on the 29K TCGA dataset.
  c. *Phikon-V2:* ViT-Large model trained on 60K WSIs from CPTAC^19^, TCGA and GTEx^20^.
  d. *UNI2 and Virchow2:* ViT-Huge models trained on approximately 300K and 3.1M proprietary WSIs, respectively
2. We evaluate these models on a diverse suite of downstream tasks that reflect real-world challenges—including tumor detection, subtype classification, and survival analysis. The benchmark suite includes both public datasets and two novel proprietary datasets, providing a comprehensive assessment across prominent and prevalent tissue types.
3. We assess model performance at both the whole slide and region-of-interest (ROI) levels, providing a comprehensive comparison across local and global tasks.

Our evaluation offers insights into the interplay between model scale, pretraining data volume, and task complexity. By highlighting these performance trade-offs, our work seeks to support researchers, clinicians, and computational pathology teams in making more informed foundation model adoption decisions, bridging the gap between technical innovation and operational impact.

## Methodology

To systematically assess the performance and trade-offs of the selected FMs, we developed an evaluation framework built around two key components: (1) a diverse suite of downstream tasks reflecting common clinical and research challenges, and (2) a standardized evaluation pipeline applied uniformly to all model. This approach ensures that our findings are both robust and directly comparable, providing a clear picture of each model’s capability.

### Downstream Tasks

To ensure practical relevance, we designed a benchmark suite where the tasks intentionally vary in several key dimensions: task type (classification vs. survival analysis), image level (whole-slide vs. region-of-interest), and tissue type. This diversity is crucial for generating meaningful insights into how different FMs perform under varied and challenging real-world conditions.

#### Whole-slide Image (WSI) Level Tasks

WSI-level tasks challenge a model to produce a single, holistic prediction for an entire slide, mirroring many real-world diagnostic and prognostic scenarios.

1. *<u>Lymph Node Metastasis Detection</u>*^25^: We use the canonical Camelyon16 benchmark to assess binary classification. The task is to classify H&E slides of sentinel lymph nodes as either containing breast cancer metastases (“tumor”) or not. The dataset consists of 398 WSIs acquired at 0.25 microns per pixel (mpp), initially partitioned into non-overlapping patches of size of 512 × 512 pixels, filtering patches having >=75% tissue region and then all resized to 224 × 224 pixels. This collection was then split into training (n=242), validation (n=27), and test (n=129), with nearly balanced classes (41 % tumor, 59 % normal).
2. *<u>Lung Cancer Subtype Classification</u>*: We also assess performance on a fine-grained histology classification task by asking models to distinguish between the two most common Non Small Cell Lung Cancer (NSCLC) subtypes: lung adenocarcinoma (LUAD) and lung squamous cell carcinoma (LUSC). For this, we use a proprietary cohort of 608 WSIs scanned at 0.5 mpp, partitioned into size of 512 × 512 pixels, filtering patches having >=75% tissue region and then resized to 224 × 224 pixels. The dataset was split into training (n=424), validation (n=91), and test (n=93) sets, with a well-balanced distribution of subtypes (59 % LUAD, 41 % LUSC).
3. *<u>Survival Analysis in Non-small Cell Lung Cancer</u>:* To evaluate prognostic capabilities, we task models with predicting overall survival (OS) directly from baseline H&E slides of NSCLC patients, where they were treated with immunotherapy and chemotherapy. This uses a proprietary clinical cohort of 286 WSIs scanned at 0.5 mpp. These images were partitioned into non-overlapping patches of size 224 × 224 pixels, filtering patches having >=75% tissue region. The dataset was split into training (n=226), validation (n=57) sets.

#### Region of Interest (ROI) Level Tasks

ROI-level tasks directly evaluate the quality of a foundation model’s patch-level embeddings. By classifying a single pre-defined region of interest (or patch), these tasks assess the core feature extraction capabilities of each FM. Unlike WSI-level analyses, this evaluation is independent of any slide aggregation model, offering a more direct measure of the embeddings’ descriptive power.

1. *<u>Breast Cancer Histology Classification</u>*^*26*^: This task tests performance on a balanced, multi-class problem requiring fine-grained morphological distinctions. The objective is to classify 400 ROI images into four distinct histological categories: normal, benign, in situ carcinoma, and invasive carcinoma. The dataset, split into training (n=320) and validation (n=80) sets, contains evenly distributed classes, with ROIs of size 2048 x 1536 pixels captured at 0.42 mpp and resized to 224 x 224 pixels.
2. *<u>Colorectal Polyp Classification</u>*^*27*^: To evaluate model robustness in a realistic scenario with significant class imbalance, we include a colorectal polyp classification task. The objective is to classify 9,536 ROIs (1812 x 1812 pixels captured at 0.44 mpp) of colorectal tissue into six different polyp subtypes. The dataset, resized to 224 x 224 pixels and split into training (n=6,270) and validation (n=2,399) sets, features a highly imbalanced class distribution, a well-known challenge in medical imaging.
3. *<u>Pan-Cancer Tissue Classification</u>*^*28*^: Finally, we use a large-scale, multi-domain benchmark to directly test the generalizability and scalability of the FM’s learned representations. The objective is to classify tissue patches into one of 31 cancer types or normal tissue categories. The massive dataset consists of 271,710 ROIs (256 x 256 pixels) extracted from The Cancer Genome Atlas (TCGA) WSIs and is split into training (n=230,850) and validation (n=40,860) sets, covering a wide spectrum of human cancers.

### Evaluation Design

Our evaluation strategy is explicitly designed to facilitate a fair and rigorous comparison across different tasks. The primary goal is to isolate the impact of the foundation model’s features. To do this, we keep all other factors consistent, ensuring that performance differences can be attributed directly to the pretrained feature extractor itself. We employ a standardized evaluation pipeline with a fixed downstream model architecture and training protocol for each task type—WSI classification, WSI survival analysis, and ROI classification. This methodology intentionally prioritizes fairness and comparability over tuning each model to its absolute peak performance on any single task.

#### Weakly Supervised Training for WSI-Level Analysis

For all WSI-level tasks, we employed a consistent weakly supervised framework where we use the frozen FM backbone to generate patch-level embeddings from center-cropped patches of size 224 x 224 pixels, with the patches normalized using the statistics specific to its corresponding FM.. These embeddings were then processed by commonly used Attention-based Multiple Instance Learning (AMIL) algorithm^29^ to produce a slide-level prediction. The specific training protocol was adapted based on the task type as described below.

##### Classification Tasks

For the Lymph Node Metastasis detection and Lung Subtype classification tasks, the AMIL model was trained using a standard cross-entropy loss. Within this setup, we used Tanh Attention module with a hidden dimension of 256. This was followed by a 3-layer classifier module, where each layer had a hidden dimension of 256, dropout rate of 0.5 and Relu activation, to finally generate final slide-level prediction. To ensure a fair comparison, a fixed configuration was used across all experiments: the model was trained for up to 100 epoch, including a 30-epoch warmup period using learning rate of 1e-4 with a OneCycleLR schedule and batch size of 1 which included all extracted patches from an entire WSI. Additionally, Early stopping based on validation performance was enabled to prevent overfitting.

##### Survival Analysis

For the NSCLC OS prediction task, the same AMIL architecture was used, but the model was trained using the negative log partial likelihood of the Cox proportional hazards model as the loss function, with a batch size of 1, while keeping rest of the configurations same as used for classification tasks. Given the limited sample size of this cohort (n=286), a 5-fold cross-validation strategy was employed to ensure a robust and stable evaluation, rather than using a single, fixed data split.

#### Linear Probing for ROI-Level Analysis

To assess the intrinsic quality and linear separability of the features from each FM, we employ a linear probing protocol for all ROI-level tasks. For our linear probing analysis, we trained an l2-regularized logistic regression model directly on the patch embeddings extracted from the frozen FM backbone. The model was optimized using an L-BFGS solver with the regularization coefficient *λ* fixed to 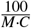, where M is the feature embedding dimension and C is the number of classes. By intentionally limiting the capacity of the downstream model, this approach ensures that performance directly reflects the “out-of-the-box” utility and quality of the learned feature space, providing a clear and efficient comparison across models.

#### Performance Metrics

To ensure a robust comparison, we evaluate model performance using a set of standard metrics appropriate for each task type.

*<u>WSI-Level Classification</u>:* For the classification tasks at the WSI level (i.e, Lymph Node Metastasis Detection and Lung Cancer Subtype Classification), we report the Area Under the Receiver Operating Characteristic Curve (AUROC).

*<u>ROI-Level Classification</u>:* For the multi-class ROI-level tasks, we report the Balanced Accuracy (BACC). This metric is robust to class imbalance as it measures the average per-class accuracy, making it a standard for this type of problem.

*<u>Survival Analysis</u>:* For the NSCLC OS prediction task, we report the mean Concordance Index (C-Index), the standard metric for evaluating the ranking performance of survival models.

All experiments were conducted using an NVIDIA L40S GPU (48 GB VRAM). The software stack included Python 3.9, PyTorch 2.5, and PyTorch Lightning 2.4.

## Results

This section presents the quantitative performance of the foundation models on WSI-level and ROI level tasks. The primary findings are summarized in Table 2 and Table 3.

**Table 1.**
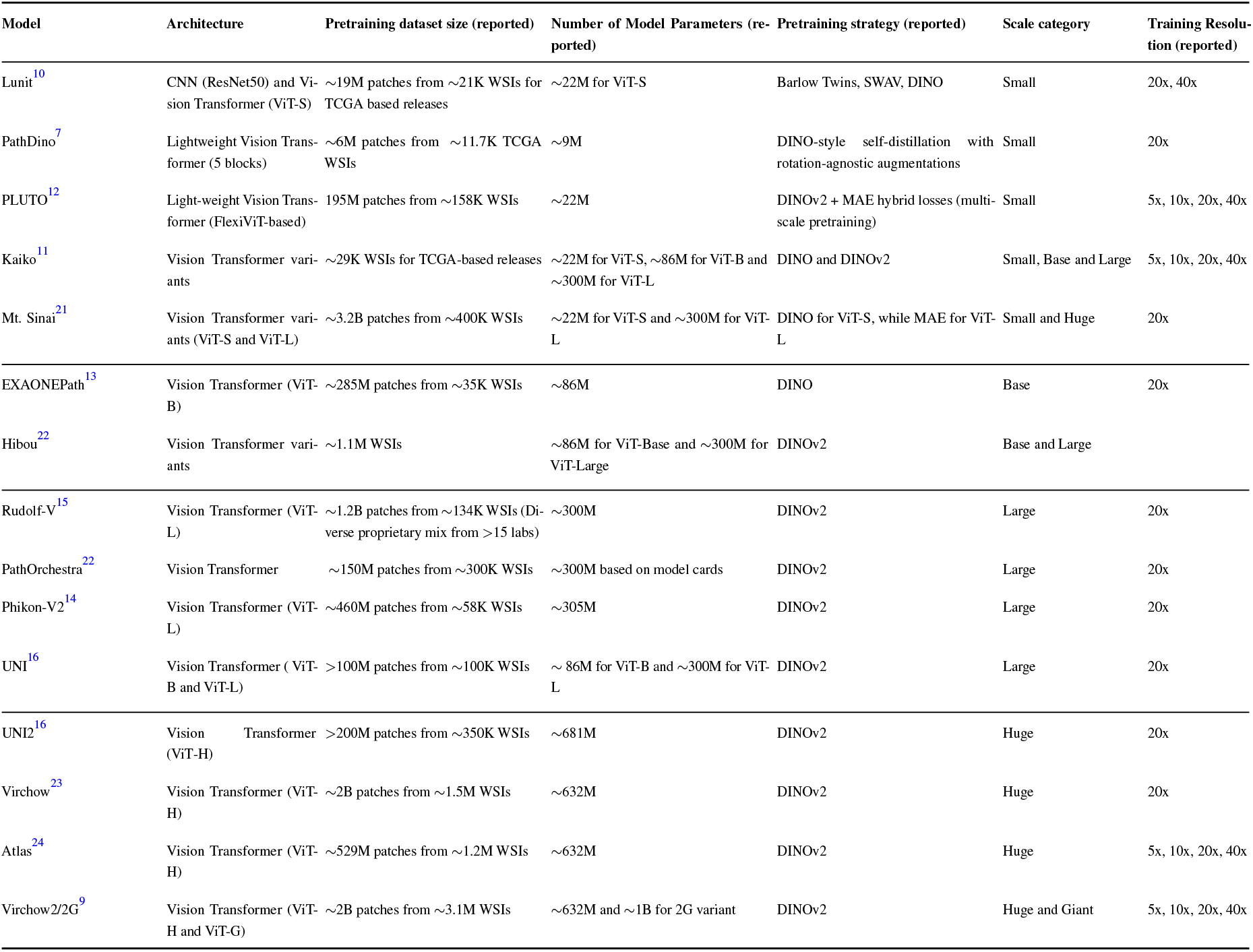
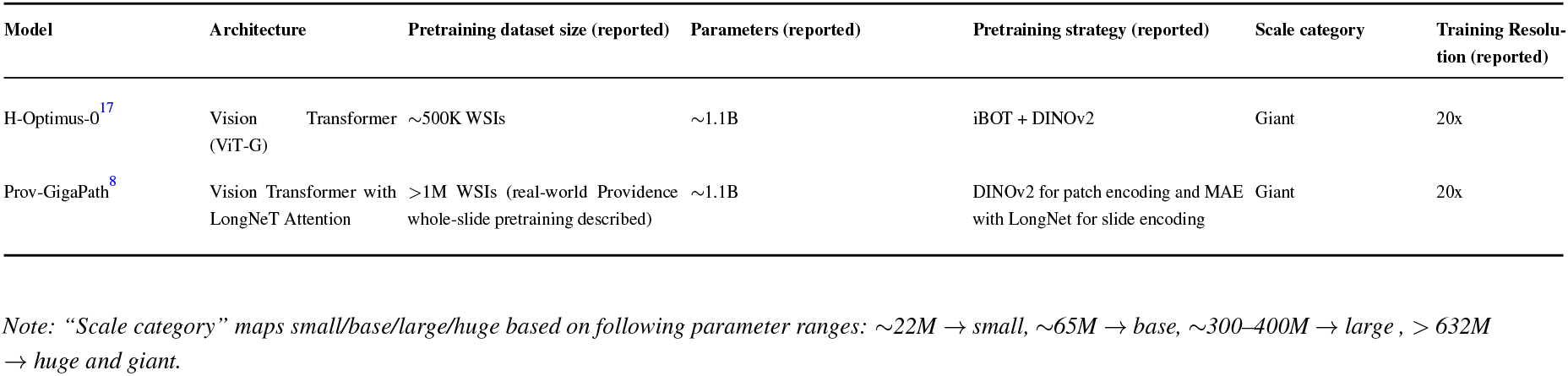
Summary of Digital Pathology Foundation Models and their characteristics.

**Table 2.**
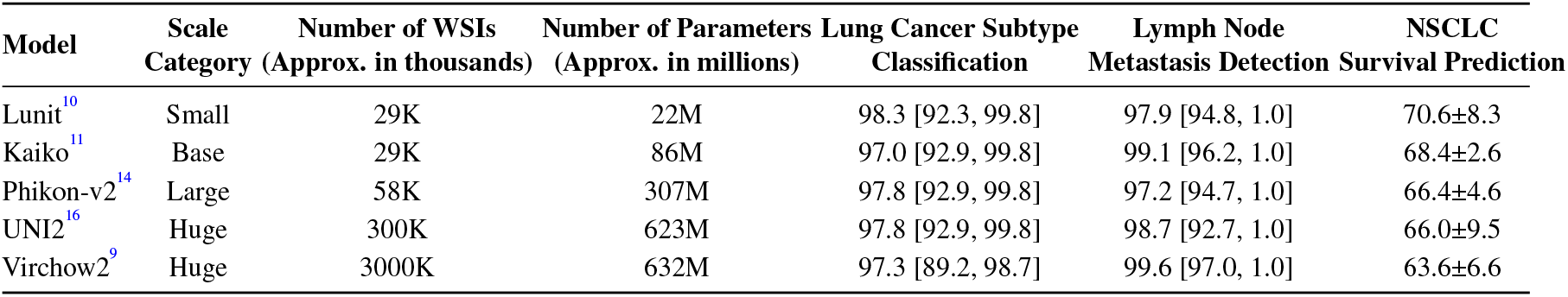
Comparative performance of selected FMs on WSI-Level tasks, measured by AUROC (alongwith confidence interval) for classification tasks (Columns 5 and 6), and C-Index (mean ± SD) for the survival analysis task (Last Column).

**Table 3.**
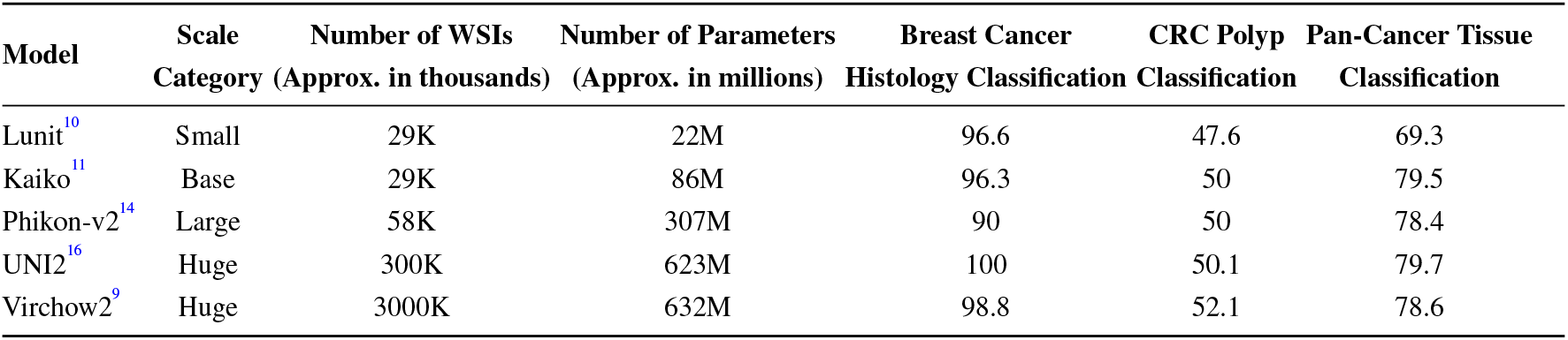
Comparative performance of selected FMs on ROI-Level tasks, measured by Balanced Accuracy(%)

### WSI-Level Analysis

Our evaluation of WSI-level tasks (Table 2) shows that no single foundation model achieves superior performance across all benchmarks. Performance varies between classification and survival analysis.

On the WSI classification tasks, performance is highly dependent on the specific benchmark dataset. For the Lymph Node Metastasis Detection task, the huge-scale Virchow2 model achieved the top AUROC of 99.6. However, the much smaller Kaiko model performed at a remarkably close level (99.1). The trend inverted on the fine-grained Lung Cancer Subtype Classification task, where the smaller Lunit model achieved the highest AUROC (98.3), followed by the larger Phikon-v2 and UNI2 models (both 97.8).

For the prognostic task of predicting overall survival in NSCLC, performance trends diverged from those observed in classification. Lunit, the smallest model, achieved the highest mean concordance index (70.6 ± 8.3) but with considerable variance. Kaiko obtained a slightly lower mean C-Index (68.4 ± 2.6) yet showed markedly greater stability. Notably, both TCGA-pretrained models outperformed the huge-scale counterparts UNI2 (66.0±9.5) and Virchow2 (63.6±6.6).

### ROI-Level Analysis

Evaluation of ROI-level tasks (Table 3) further demonstrates task-specific performance differences among models.

The analysis is dominated by the exceptional performance of the UNI2 model, which achieved a perfect Balanced Accuracy score (1.0) on the Breast Cancer Histology Classification (BACH) task and also secured the leading score on the broad Pan-Cancer Tissue Classification task (79.7).

The advantages of massive-scale, however, were apparent in the most challenging benchmark. The VIRCHOW2 model achieved the top score on the CRC Polyp Classification task (52.1). Among the smaller models, Kaiko (50.0) demonstrated strong generalization, outperforming Lunit (47.6) on the CRC and Pan-Cancer tasks.

## Discussion

### Is Model Scale a Universal Predictor of Performance?

Our benchmarks demonstrate that the effect of model scale is far from uniform across tasks. At the WSI-level, while Huge-scale model (i.e. Virchow2) achieved the highest AUROC (99.6) in broad screening challenges like lymph node metastasis detection, at the same time, a base-scale model (i.e. Kaiko) demonstrated highly competitive performance. Furthermore, In fine-grained classification, the smallest model, Lunit, achieved the strongest performance (98.3 AUROC on lung cancer subtype classification), further demonstrating that larger model scale was not universally beneficial for certain tasks. This pattern was even more pronounced in the survival analysis task, where the ViT-small and ViT-base models (i.e. Lunit and Kaiko), outperformed their huge-scale counterparts (i.e. UNI2 and Virchow2). Several factors may explain this phenomenon. First, the simple AMIL aggregator used in our standardized pipeline may act as an information bottleneck, potentially struggling to effectively process the higher-dimensional embeddings from larger models. Second, and perhaps more critically, is the nature of the task itself. Prognostic tasks like survival analysis often depend on a mix of morphological and non-morphological factors (e.g., patient demographics, molecular status, tumor stage). It is plausible that larger models, with their vast capacity, overfit to the purely morphological features in the training data, failing to generalize as well as smaller models that learn more robust, less complex representations. To summarize, these findings suggest a crucial insight: simply increasing model scale does not guarantee superior performance. While larger models may excel at broad morphological classification, their advantage diminishes and can even reverse in more specialized or complex prognostic tasks where outcomes are not solely determined by visual patterns

### Do Smaller Models Trained on Limited Datasets Suffice?

Our analysis provides compelling evidence that models with a smaller scale, trained on moderately sized datasets, can achieve highly competitive and sometimes superior downstream performance compared with large/huge-scale models. Kaiko (base-scale, 29k WSIs) exemplifies this: it achieved better performance against Phikon-v2 (large-scale, 58K WSIs), while still being competitive against UNI2 (huge-scale, 300K WSIs) on Pan-Cancer classification, demonstrated comparable performance to Virchow2 (huge-scale, 3.1M WSIs) on metastasis detection, and delivered the most stable survival predictions of all models evaluated. Similarly, Lunit, despite being the smallest in model and data scale, achieved the highest AUROC for lung subtype classification and the top mean C-index, albeit with greater variance. These findings challenge the common notion that a massive scale of pretraining data as well as model is always necessary. They suggest that smaller models can be highly effective, particularly for specialized or prognostic tasks, when pretrained on high-quality, curated datasets with sufficient diversity. For many practical applications, such models represent an attractive trade-off between performance, robustness, and computational cost.

### Is Model Superiority Task-Specific or Generalizable?

The cross-task comparisons show that model performance is strongly task-dependent. No single foundation model consistently dominated across all benchmarks. For instance, Virchow2 excelled at metastasis detection and CRC polyp classification, while UNI2 was the strongest for ROI-level breast cancer subtyping and Pan-Cancer tissue classification. Notably, Kaiko consistently delivered competitive accuracy across both WSI- and ROI-level tasks while also demonstrating remarkable robustness in survival prediction. These results underscore that performance gains do not automatically generalize from one domain to another, and that the design choices beyond sheer scale are critical. The observed performance differences may be linked to the model’s pretraining strategy. For example, UNI2’s strength in diverse, fine-grained ROI classification could be attributed to its pretraining on a large, multi-institutional dataset that fosters robust feature learning^16^. Conversely, Virchow2’s success in WSI-level screening aligns with its pretraining on data from multiple magnifications (5x, 10x and 20x), a strategy designed to explicitly teach the model to integrate cellular detail with broader tissue context^23^. This suggests a “data-task resonance”, highlighting the importance of aligning pretraining data and model design with the intended application, rather than assuming global superiority of any single model.

### Balancing Efficiency and Cost in Real-World Deployment

Beyond performance, deployment considerations highlight a further dimension of trade-offs. Training cost is largely a one-time expense shouldered by developers, but inference cost of running models on patient slides daily determines long-term clinical viability. Operational efficiency is shaped by hardware constraints, batch size, and throughput, and these factors can dramatically influence cost per slide in practice. For example, if we consider an inference-optimized NVIDIA L4 GPU, smaller models such as Lunit (~22M parameters) would offer minimal memory footprint, enabling large batch sizes and low per slide cost. By contrast, massive-scale models like UNI2 or Virchow2 (~632M parameters) consume significantly more memory, limiting throughput and increasing operational expenses. In this context, Kaiko provides a compelling middle ground, offering competitive accuracy with greater efficiency and scalability than the largest models. Quantifying such cost-performance trade-offs is essential for responsible adoption of digital pathology FMs. Incorporating efficiency and cost perspectives alongside accuracy provides a holistic framework for model selection, ensuring that technical advances translate into sustainable and clinically deployable solutions.

## Conclusion

Our comparative analysis demonstrates that selecting a foundation model in digital pathology is a strategic decision that must be guided by the specific requirements of the intended application. The optimal choice is rarely the largest or most data-intensive model; rather, it is the one whose model scale and image resolution strategy align best with the task. While our comparisons cannot fully disentangle the effect of training data composition, the differences observed between models pretrained on proprietary versus public datasets suggest that data sources may also play a critical role, particularly in prognostic tasks.

Notably, smaller models can outperform massive models in complex prognostic tasks such as survival analysis, while mid-sized models like Kaiko strike a practical balance between efficiency, accuracy, and stability. Beyond raw performance, reliability and consistency across runs emerge as equally important, particularly in prognostic applications where variance can undermine clinical trust.

Equally, practical considerations of computational efficiency and cost play a central role in real-world adoption. While massive models may be appealing when accuracy is the sole criterion, their high resource demands can make them less suitable for routine diagnostic workflows. In contrast, models that deliver competitive accuracy with lower inference costs offer a more sustainable path for integration into production settings. Even modest efficiency gains can translate into substantial operational savings when scaled across large patient cohorts.

While this study provides a practical framework for understanding the trade-offs of foundation model scale, it is important to acknowledge its limitations. First, our selection includes representative, but not exhaustive models for each scale tier. We posit that our findings offer a valuable general guideline, though exceptions will undoubtedly exist. Second, while the inclusion of both public and proprietary datasets provides novel insights, every dataset has inherent constraints, such as limited sample size or specific clinical contexts (e.g., the internal clinical trial cohort includes only late-stage NSCLC patients). Therefore, our results should be interpreted as an informative case study rather than a universally definitive benchmark. Finally, due to computational constraints, our analysis prioritized model diversity across different development teams over an exhaustive comparison of all scaled versions within a single model family (e.g., UNI-Large vs. UNI2-Huge). This design choice allowed for a broader survey of the landscape, but future work should include rigorous intra-family comparisons to further dissect the precise impact of scale.

In summary, our findings suggest that effective deployment requires a holistic approach, which requires balancing task-specific alignment, performance stability, computational efficiency, and cost sustainability. Such a framework ensures not only robust benchmark performance but also genuine clinical applicability, paving the way for foundation models to be responsibly and impactfully adopted in digital pathology.

## Data Availability Statement

The dataset used for Lung Cancer Subtype Classification and NSCLC Survival Prediction tasks are proprietary and cannot be made publicly available. The dataset for Lymph Node Metastasis Detection and all datasets part of the ROI-Level Analysis are publicly available and can be found as per below:

- Lymph Node Metastasis Detection: https://camelyon16.grand-challenge.org/Data/
- Breast Cancer Histology Classification: https://zenodo.org/records/3632035
- CRC Polyp Classification: https://zenodo.org/records/4643645
- Pan-Cancer Tissue Classification: https://zenodo.org/records/3373439

## Acknowledgements

We extend our sincere gratitude to our collaborators at the Genentech gCS DP team: Dan Ruderman, Cyrus Manuel, Evan Liu, and Clinton Mielke. Their partnership was invaluable, contributing significantly to benchmarking discussions, compilation of various foundation model metadata, and enriching this project through many insightful discussions.

## Author contributions statement

D.M.: Conceptualization, Methodology, Data curation, Investigation, Software, Writing – review & editing. N.S.: Conceptualization, Methodology, Data curation, Investigation, Writing – original draft & review & editing. X.L.: Conceptualization, Methodology, Investigation, Visualization, Writing coordination, Writing – original draft & review & editing. Y.N.: Conceptualization, Methodology, Investigation, Writing – review & editing. R.V.: Conceptualization, Project Supervision, Resources, Writing – review & editing. U.K.: Conceptualization, Methodology, Investigation, Funding acquisition, Project Supervision, Writing – review & editing.

